# Phase-amplitude markers of synchrony and noise: A resting-state and TMS-EEG study of schizophrenia

**DOI:** 10.1101/740621

**Authors:** D Freche, J Naim-Feil, S Hess, A Peled, A Grinshpoon, E Moses, N Levit-Binnun

**Affiliations:** Sagol Center of Brain and Mind, Ivcher School of Psychology, Interdisciplinary Center (IDC), Herzliya, Israel; Department of Physics of Complex Systems, Weizmann Institute of Science, Israel; Turner Institute for Brain and Mental Health, School of Psychological Sciences; Monash Biomedical Imaging, Monash University, Clayton, Australia; Geha Mental Health Center, Petah Tikvah, Israel; Sackler Faculty of Medicine, Tel Aviv University, Ramat Aviv, Israel; Rappaport Faculty of Medicine, Technion; Institute for Psychiatric Studies, Shaar Menashe Mental Health Center, Israel

## Abstract

The electroencephalogram (EEG) of schizophrenia patients is known to exhibit a reduction of signal-to-noise ratio and of phase locking, as well as a facilitation of excitability, in response to a variety of external stimuli. Here we demonstrate these effects in transcranial magnetic stimulation (TMS)-evoked potentials and in the resting-state EEG. To ensure veracity we used three weekly sessions and analyzed both resting state and TMS-EEG data. For the TMS responses our analysis verifies known results. For the resting state we introduce the methodology of mean-normalized variation to the EEG analysis (quartile-based coefficient of variation), which allows for a comparison of narrow-band EEG amplitude fluctuations to narrow-band Gaussian noise. This reveals that amplitude fluctuations in the delta, alpha and beta bands of healthy controls are different from those in schizophrenia patients, on time scales of tens of seconds. We conclude that the EEG-measured cortical activity patterns of schizophrenia patients are more similar to noise, both in alpha and beta resting state and in TMS responses. Our results suggest that the ability of neuronal populations to form stable, locally and temporally correlated activity is reduced in schizophrenia, a conclusion that is in accord with previous experiments on TMS-EEG and on resting-state EEG.

## 1 Introduction

Schizophrenia is a mental disease with a complex pathology that dynamically evolves across the adult lifespan (Krystal et al., 2017). Several decades of research in schizophrenia have provided extensive evidence to the pervasive, globally occurring structural, physiological, functional and genetic abnormalities that characterizes the disorder. Abnormalities of brain tissue structure from the scale of synapses to whole brain regions have been described, with connectivity degradation occurring both locally and long-range. Correspondingly, a range of profound functional abnormalities were observed in activity, from cell assemblies to large-scale networks (Uhlhaas and Singer, 2010; Krystal et al., 2017).

One way to probe the functional abnormalities in schizophrenia is by studying dynamics of brain activity using EEG in stimulus-response paradigms. Generally, the main alterations of evoked responses in schizophrenia concern reductions in amplitudes and in phase locking (Winterer et al., 2000, 2004; Spencer et al., 2003; Frantseva et al., 2012; Shin et al., 2015), as well as facilitated excitation in late-stage response (Radhu et al., 2015; Frantseva et al., 2012; Rogasch et al., 2014), increased inter-trial variability (Winterer et al., 2000, 2004), and altered spectral power (Winterer et al., 2000; Frantseva et al., 2012).

A hypothesis to explain these findings, first formulated by Winterer et al. (2000), suggests the presence of inherent noise in brain activity. This manifests itself as stimulus-unrelated background activity that interferes with stimulus-evoked responses and thus leads to increased inter-trial variability. Further work by Rolls et al. (2008) conceptualized this hypothesis within a dynamical systems framework, with altered stability properties of attractors. Krystal et al. (2017) suggested that although the large inter-trial variability observed in schizophrenia can result from random background activity that degrades regular signals, it may also lead in itself to the creation of an aberrant signal. This suggests that not only evoked activity should be affected by such noise, but that the resting state activity in schizophrenia should exhibit correlates of abnormal activity as well. Indeed, various alterations of the resting-state EEG in schizophrenia have been reported. General findings include alterations of power spectra (Uhlhaas and Singer, 2010) and abnormalities of spatial activity in analyses of microstates (Rieger et al., 2016; Michel and Koenig, 2018). Notably, temporal activity alterations in long-range time correlations (LRTC) (Nikulin et al., 2012; Sun et al., 2014; Moran et al., 2019) were found, demonstrating that brain activity in patients shows reduced temporal auto-correlations on the timescale of tens of seconds, as well as reduced amplitude excursions (Sun et al., 2014).

This hypothesis that noise and its regulation is at the heart of alterations of functional activity in schizophrenia has been suggested as a low-level mechanism to explain both physiological and behavioral findings (Northoff and Duncan, 2016; Winterer et al., 2000; Rolls et al., 2008). Alterations in global activity can shape the processing of stimulus-induced activity and may provide the basis for the reported changes in cognitive, affective and social functions (Northoff and Duncan, 2016). Ultimately, these alterations may underlie the unstable symptoms and symptom severity characteristic of the disorder (Habtewold et al., 2019; Bota et al., 2011). Despite the apparent popularity of the noise hypothesis, the fundamental question regarding the nature of this postulated noise remains to be answered. This regards its manifestations at the molecular, synaptic, or cellular level, as well as its properties, such as more deterministic or more random characteristics, permitting different interpretations of variability in brain activity (Faisal et al., 2008).

Here, we formulate the noise hypothesis in the following way: The presence of noise is a characteristic of brain activity in schizophrenia, which reduces the capacity of the brain to achieve the formation of spatially and temporally stable correlated activity in neuronal populations. In this formulation, the hypothesis applies to brain activity in response to a stimulus as well as in resting state. This makes it possible to view the TMS-EEG as well as the resting-state results in a unified context.

We evaluated the strength of these correlations in population activity using markers derived from resting-state and TMS-EEG repeated-sessions experiments with schizophrenia patients and healthy controls. In TMS-EEG, we evaluated the signal-to-noise ratio, phase locking to stimulus, and post-stimulus excitability suppression. In resting-state EEG, we analyzed how amplitude averages and amplitude fluctuations compare to random noise. Generally, we find a reduction of these markers in the schizophrenia group. We suggest the interpretation that the ability to create stable spatiotemporal correlations of brain activity is reduced, consistent with the noise hypothesis.

## 2 Materials and methods

The study was approved by the Shaar Menashe Mental Health Center Institutional Review Board. All participants were reimbursed the equivalence of 35 US dollars for participating in each EEG recording sessions.

### 2.1 Experiments

We record TMS-evoked and resting-state EEG activity in schizophrenia patients and healthy controls, with three repetitions for each participant. Evaluating both groups three times with one week between each session strengthens the reliability of the results (see also Kerwin et al. (2018)), as cortical responses may vary with changes in symptoms (Naim-Feil et al., 2019). An advantage of TMS-EEG is that it does not require active participation and attention of the schizophrenia patients (Daskalakis et al., 2002; Rogasch et al., 2014).

### 2.2 Subjects

Thirty healthy controls without any history of psychiatric illness, drug abuse or head injury were recruited through local advertisements. Three subjects were excluded because not all three sessions were attended, yielding 27 datasets for control.

Thirty in-unit schizophrenia patients at Shaar Menashe Mental Health Center qualified for the study according to DSM IV-TR Schizophrenia (DSM, 2000). The patients did not have any history of neuromodulation, except one patient who had received ECT more than four years before the study. Nine subjects were excluded because not all three sessions were attended, yielding 21 datasets of schizophrenia patients. Each patient was evaluated by a trained psychiatrist using the Scale for the Assessment of Positive Symptoms (SAPS) and the Scale for the Assessment of Negative Symptoms (SANS) to quantify clinical symptoms of schizophrenia (Andreasen, 1984, 1989).

All subjects completed a general demographic questionnaire and TMS safety questionnaire (Rossi et al., 2011). For demographic and clinical data, see Table 1. Each recording session was supervised by a trained psychiatrist, who assured that epileptic activity was absent in the EEG of all subjects.

**Table 1:**
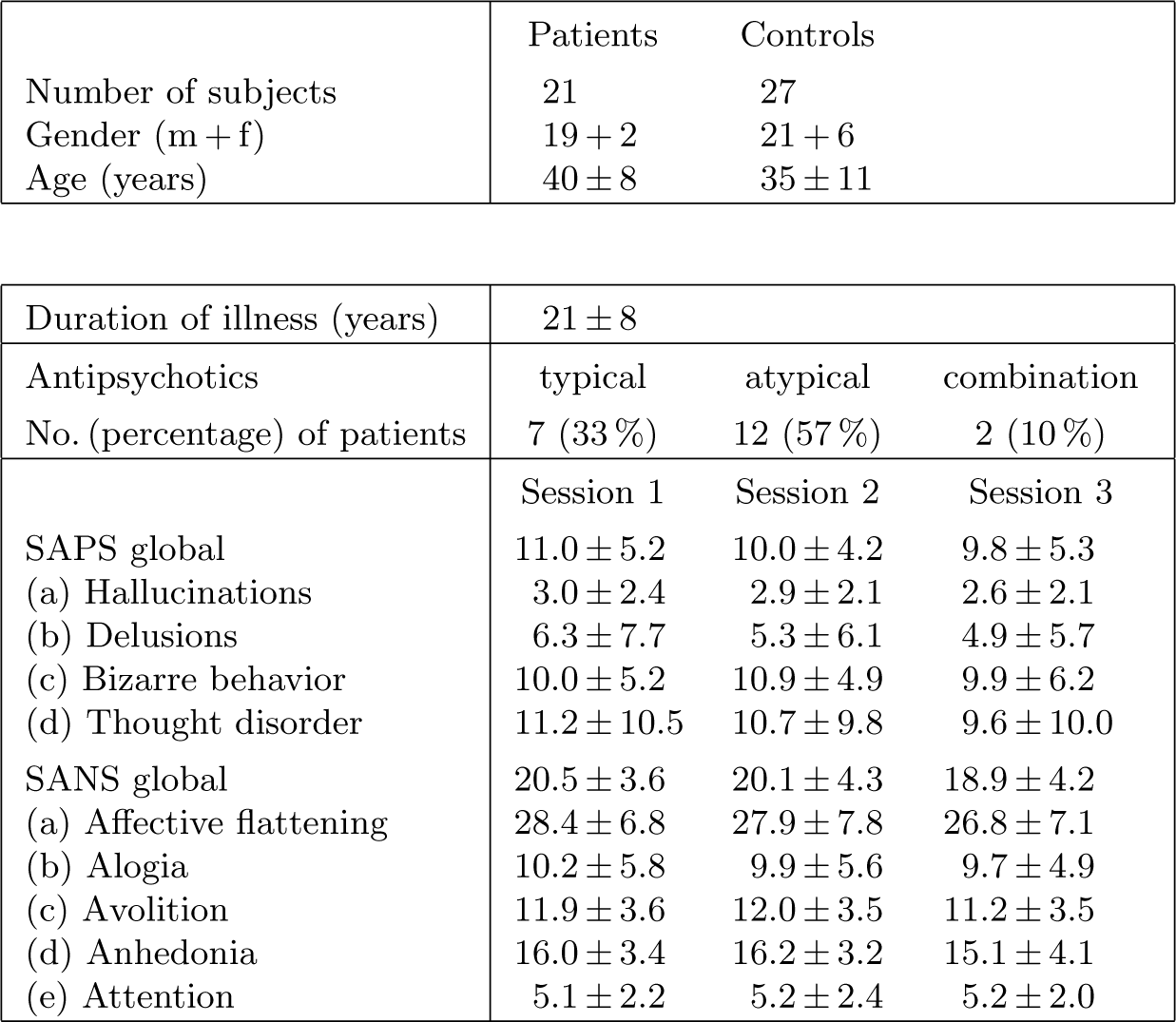
Demographic and clinical data.

|  | Patients | Controls |
| --- | --- | --- |
| Number of subjects | 21 | 27 |
| Gender (m + f) | 19 + 2 | 21 + 6 |
| Age (years) | 40 ± 8 | 35 ± 11 |

Table 1: *Demographic and clinical data.*
|  |  |  |  |
| --- | --- | --- | --- |
| Duration of illness (years) | 21 ± 8 |  |  |
| Antipsychotics | typical | atypical | combination |
| No. (percentage) of patients | 7 (33 %) | 12 (57 %) | 2 (10 %) |
|  | Session 1 | Session 2 | Session 3 |
| SAPS global | 11.0 ± 5.2 | 10.0 ± 4.2 | 9.8 ± 5.3 |
| (a) Hallucinations | 3.0 ± 2.4 | 2.9 ± 2.1 | 2.6 ± 2.1 |
| (b) Delusions | 6.3 ± 7.7 | 5.3 ± 6.1 | 4.9 ± 5.7 |
| (c) Bizarre behavior | 10.0 ± 5.2 | 10.9 ± 4.9 | 9.9 ± 6.2 |
| (d) Thought disorder | 11.2 ± 10.5 | 10.7 ± 9.8 | 9.6 ± 10.0 |
| SANS global | 20.5 ± 3.6 | 20.1 ± 4.3 | 18.9 ± 4.2 |
| (a) Affective flattening | 28.4 ± 6.8 | 27.9 ± 7.8 | 26.8 ± 7.1 |
| (b) Alogia | 10.2 ± 5.8 | 9.9 ± 5.6 | 9.7 ± 4.9 |
| (c) Avolition | 11.9 ± 3.6 | 12.0 ± 3.5 | 11.2 ± 3.5 |
| (d) Anhedonia | 16.0 ± 3.4 | 16.2 ± 3.2 | 15.1 ± 4.1 |
| (e) Attention | 5.1 ± 2.2 | 5.2 ± 2.4 | 5.2 ± 2.0 |

### 2.3 EEG recording

EEG was recorded from 64 electrodes using a Biosemi ActiveTwo amplifier (BioSemi, Netherlands) at a sampling rate of 1024 Hz. TMS was applied with a Magstim Rapid (Magstim, UK) and a figure-of-eight coil at a fixed strength of 80 percent maximal stimulator output. There were three recording sessions. Each session was recorded at the same time of day with one week between the sessions. In each session, there were two stimulation sites (above frontal and central-parietal electrodes, FC2 and either FC2 or CP2). At each of these sites, a real stimulation measurement was always coupled to a sham stimulation (by placing the TMS coil perpendicularly to the stimulation site). Each such measurement (e.g., real or sham stimulation over FC2) comprised of around 100 single TMS pulses delivered manually at ∼3 s inter-pulse intervals. Each of the three stimulation sessions was preceded by a resting-state recording (duration mean±std: 260 s±60 s) in which subjects were requested to keep their eyes open. Here we report comparative analysis only for the real stimulation, as the sham stimulation is not relevant to the questions of this paper. Results are also only analyzed for stimulation over the FC2 electrode, as this stimulation site had the most complete data sets.

To maintain consistency between sessions, several measures were undertaken. First, the same EEG cap was used for all sessions with the same cap location as manually tape-measured. Second, a custom-built mobile, manually adjustable stand for the TMS coil was used in order to keep the coil position and orientation with respect to the FC2 electrode across sessions. Third, sessions were conducted at the same time of day and subjects were requested to maintain their sleep, coffee, and cigarette intake (which was monitored) constant for the days of the experiment. Furthermore, subjects listened to white noise through headphones during the sessions to reduce auditory distraction (Paus et al., 2001; Fitzgerald et al., 2008).

### 2.4 EEG data processing

Resting-state EEG and TMS-EEG data were processed offline in several steps. In the TMS-EEG datasets, two additional pre-processing steps were applied. First, the procedure of Freche et al. (2018) was used to remove the TMS-induced discharge artifacts, and subsequently, the data of 25 ms duration following the TMS-pulse application, which could not be easily reconstructed at this sampling rate, were linearly interpolated. Afterwards, all datasets were high-pass filtered at 1 Hz and low-pass filtered at 150 Hz followed by a notch filter at 50 Hz using FIR filters by applying EEGLab’s eegfiltnew function. Bad electrodes were removed from any further analysis following visual inspection. Remaining artifacts due to muscle activity, eye blinks, etc. were removed using Independent Component Analysis (ICA). The TMS-EEG data were subsequently epoched into trials for a time window of 1 s before and 2 s after each pulse. Finally, EEG signals were re-referenced by subtracting the signal average. In resting-state EEG datasets, only the first 182 s duration was used for further analysis. After further time-frequency decomposition (see Section Data Analysis below, the 1 s at the beginning and at the end were removed to accommodate for edge effects. In the case of spectral power density computations (see Section Data Analysis below), the beginning and ending seconds were also discarded to ensure the same length, even though there were no edge effects. Six datasets (5 sessions of different controls, 1 session of one patient) had a duration below 182 s and were therefore excluded from the resting-state EEG data analysis.

Care was taken during the data acquisition to ensure that the resting-state EEG recordings did not have any bad electrodes. However, in the TMS-EEG recordings, bad electrodes did occur, and in these datasets bad electrodes had to be removed, on average 1.3 electrodes in the controls group and 6.3 electrodes in the patients group, per dataset. About 50 % of the electrodes deleted were the FC2 electrode (stimulation site) or its nearest-neighbor electrodes, which were randomly prone to occasionally touching the coil.

### 2.5 Data analysis

#### 2.5.1 Time-frequency decomposition, phase locking value (PLV), relative power, power spectral density

Time-frequency decompositions were computed using Morlet wavelets in the range of 2 Hz to 45 Hz with 2 cycles at the lowest and 15 cycles at the highest frequency in logarithmically spaced steps (cf. Bertrand et al. (2000)).

The PLV was computed from the phase of the time-frequency for each frequency and time point (Tallon-Baudry et al., 1996). It ranges between 0 and 1, where larger values indicate stronger phase locking between trials. The PLV was used to compare phases after the TMS pulse across trials.

Relative power was derived for each frequency, dividing by mean baseline power for baseline-normalization, and then converted to decibels. The baseline power was obtained as the mean power in the pre-stimulus window of 300 ms to 700 ms, averaged over all trials. This window falls outside the 2*σ*_*t*_-wide zone of influence of the trial boundaries and of the stimulation artifact, where *σ*_*t*_ is the (frequency-dependent) standard deviation in time of the Morlet wavelet (cf. Bertrand et al. (2000)).

For statistical analyses of the TMS-EEG data, time-frequency data were further processed in two steps. First, to take the same number of trials for every subject, only the first 70 trials surviving artifact removal were considered. Second, smaller aggregates were obtained, for every electrode, by down-sampling in time to 128 Hz and in frequency by averaging over the frequency bands, i.e. the delta (1-4 Hz), theta (4-8 Hz), alpha (8-12 Hz), beta (12-30 Hz), and gamma bands (30-45 Hz). In this paper, we call such an aggregate a time-frequency-electrode pixel, which is hence indexed by a single (downsampled) time point, frequency band, and electrode. Furthermore, we call a quantity a time-frequency pixel if it is specified by a single time point and a frequency band.

The power spectral density (PSD) was obtained for each electrode in the resting-state EEG by first computing the Fourier spectrum in a moving window of 2 s duration and 1 s overlap, and then taking the median of these windows.

The alpha peak location for each subject was obtained in the resting-state EEG by manual inspection of the average of the power spectral densities of all electrodes. For all three sessions of 1 control subject and one session of 1 patient, an alpha peak could not be detected, and therefore these subjects were not accounted for the peak location analysis.

#### 2.5.2 Quartile-based coefficient of variation (qbCV)

According to the common view of EEG, there are two types of components that constitute the measured activity (Sauseng et al., 2007). One originates from spontaneous, ongoing activity, and is also called background activity. The other originates from short-lived and locally correlated bursts of synchronized activity such as event-related potentials. The first can be regarded as background noise and the second as intervals of modulated oscillatory signals.

Here we use a simple marker that can indicate whether modulated oscillations are present in a white noise signal. A common measure for the spread of a distribution based on its percentiles is the quartile-based coefficient of variation (qbCV). See Arachchige et al. (2019) for a discussion of its statistical properties. Defining *Q*_1_ as the percentile below which 25 percent of the distribution resides and *Q*_2_, *Q*_3_ the corresponding values for the 50-th and 75-th percentile, respectively, the qbCV is defined by (*Q*_3_ *− Q*_1_)*/Q*_2_. It is well-known that narrow-band-filtering of pure white noise yields a Rayleigh-distributed instantaneous amplitude (McClaning, 2012) whose scale parameter is the amplitude of the underlying white noise. This distribution has the property that its quantile function linearly depends on its scale parameter. This implies that the amplitude average is proportional to its fluctuation average, and hence that its qbCV is independent of the amplitude of the underlying white noise. Furthermore, its qbCV has a known analytic expression with *qbCV*_*Rayleigh*_ ≈ 0.77. Thus, a signal with a qbCV different from *qbCV*_*Rayleigh*_ implies the presence of non-noise components such as oscillatory signals.

For example, a pure-noise narrow-band signal with an additional constant oscillation at the mid-band frequency would increase the mean of the amplitude and hence the qbCV would be smaller than *qbCV*_*Rayleigh*_. Another example is a pure-noise narrow-band signal with additional intervals of oscillatory bursts. This will increase the variation of the signal amplitude, leading to an increase in qbCV relative to *qbCV*_*Rayleigh*_.

For the resting-state EEG analysis of the section on amplitude fluctuations below, the quartiles *Q*_*i*_ (*i* = 1, *dotso*, 3) were derived for every frequency from 1 to 45 Hz from the time-frequency decomposition. For each electrode, a *Q*_*i*_ was computed in a moving window of 40 s duration and 20 s overlap, and then the median of all windows was taken. Finally, the qbCVs were computed from these medians.

### 2.6 Statistical analysis

Each statistical analysis was first performed at each electrode separately. The purpose of the single-electrode analyses was to evaluate phase and amplitude markers for correlated neural activity locally, i.e. at the single-electrode level, in the context of the noise hypothesis as stated in the introduction. Furthermore, it allows to observe that the topographical distribution of all of the significantly different electrodes does not occur randomly on the head but rather follows physiologically reasonable patterns such as hemispheric symmetry or proximity to the TMS stimulation site. Following the electrode-wise tests, we also accounted for multiple comparisons using an FDR-controlling procedure. Because the various quantities we tested may be correlated in various (unknown) ways, we used the Binyamini-Yekutieli method for correction. The significance threshold was set to *α* = 5 %.

Statistical comparisons of relative power (for Fig. 2) and PLV (for Fig. 3) were performed based on a mass univariate analysis approach (Groppe et al., 2011). First, group differences were tested for all time-frequency-electrode pixels separately. Excluded from this analysis were all pixels which were outside the *σ*_*t*_-wide zone of influence of the trial boundaries and of the stimulation artifact (shown in gray in Figs. 2A, 3B). We used the following statistical two-level model. The first level describes the session-specific effects, and the second the subject-specific effects. For a quantity *Y*_*ij*_ of session *i* and subject *j*,

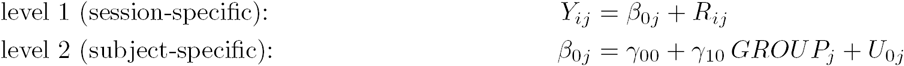

**Figure 1:**
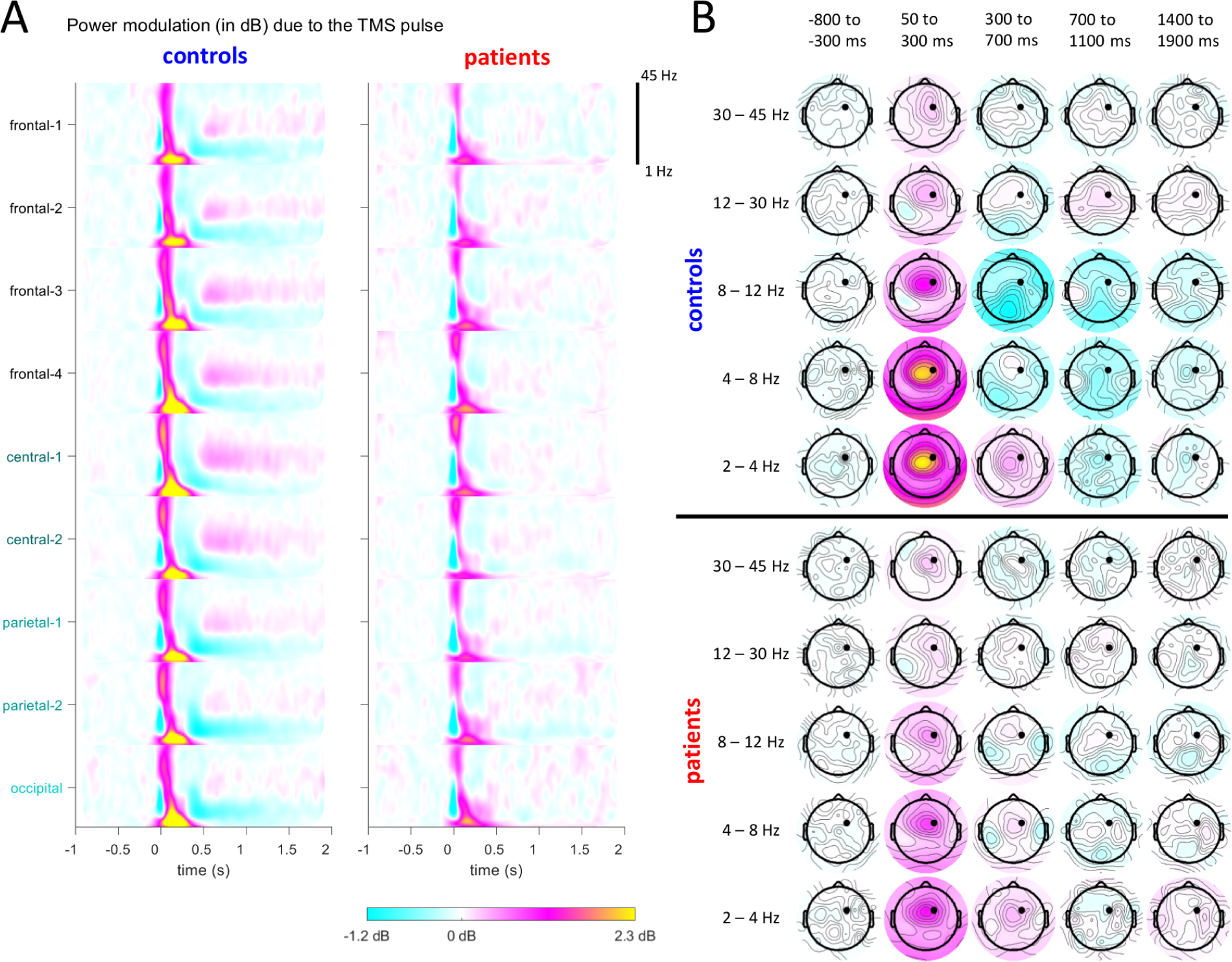
**A.** Time-frequency decompositions for electrode aggregates (averaged across trials). TMS was applied at time 0. The smearing in the time-frequency plot around time zero is due to the artifact removal process which introduces a gap of 25 ms in the data following the pulse onset. **B.** Topographical plots for different time-frequency windows (averaged over windows and across trials). The stimulation site FC2 is marked by a black dot.

**Figure 2:**
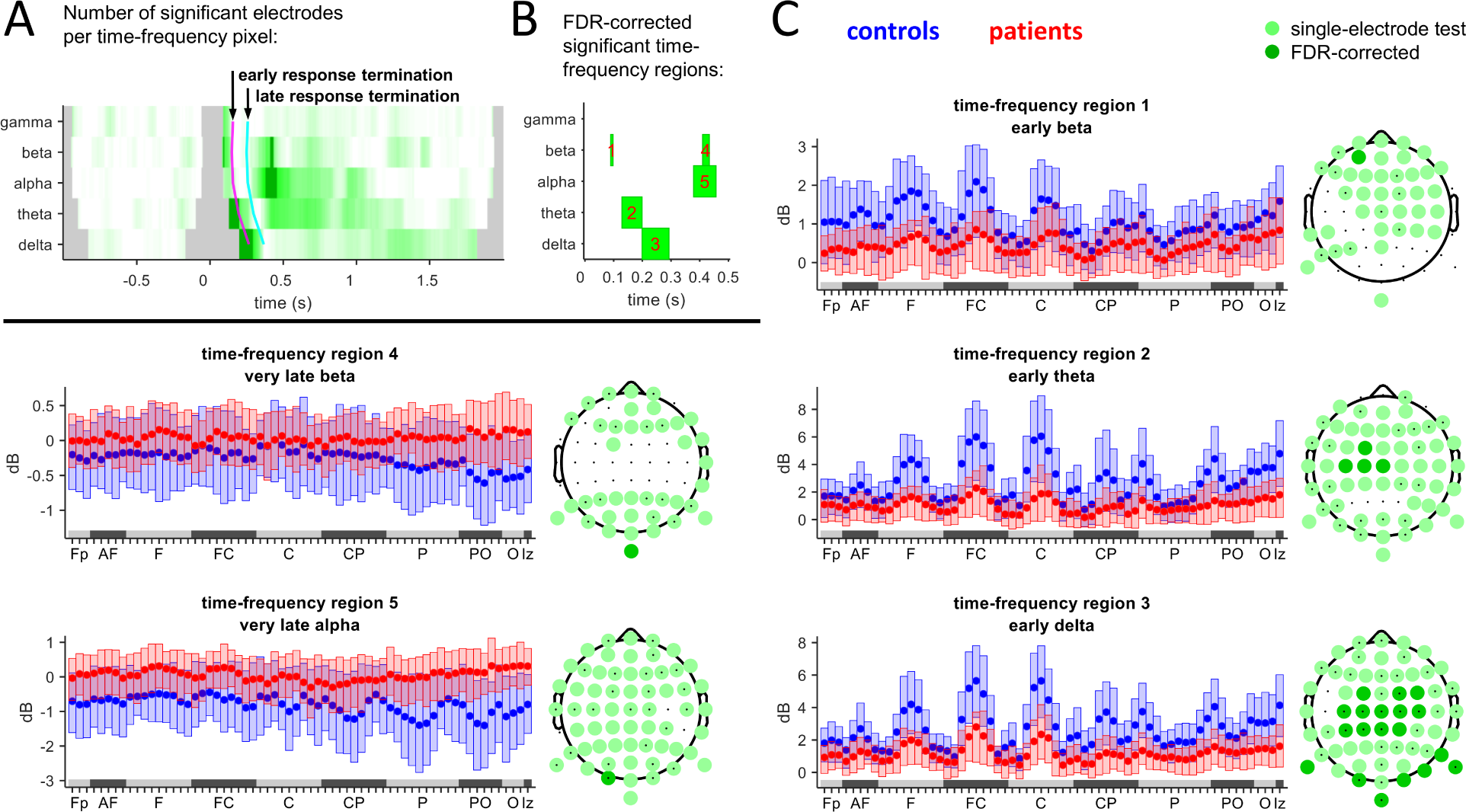
**A.** For the power values at each time-frequency-electrode pixel, a statistical test was performed (p ≤ α=5 %). Shown is the number of electrodes that are significant at a time-frequency pixel. More intense green means larger number of significant electrodes. Dark green: Time-frequency pixels selected from those FDR-corrected time-frequency pixels for which there is at last one significant time-frequency-electrode pixel. Gray: excluded due to boundary and artifact interference. Early response: time span until 100 ms (magenta line), late: 100 ms to 200 ms (cyan line). **B.** Time-frequency regions obtained from grouping the dark green time-frequency pixels by frequency band and by response stage. **C.** Boxplots of the power values of each time-frequency region for each electrode (x-axis: 64 electrodes aggregated by groups. Order inside a group is band-wise from the right to left hemisphere.) Statistically significant group differences are shown as topographical plots.

**Figure 3:**
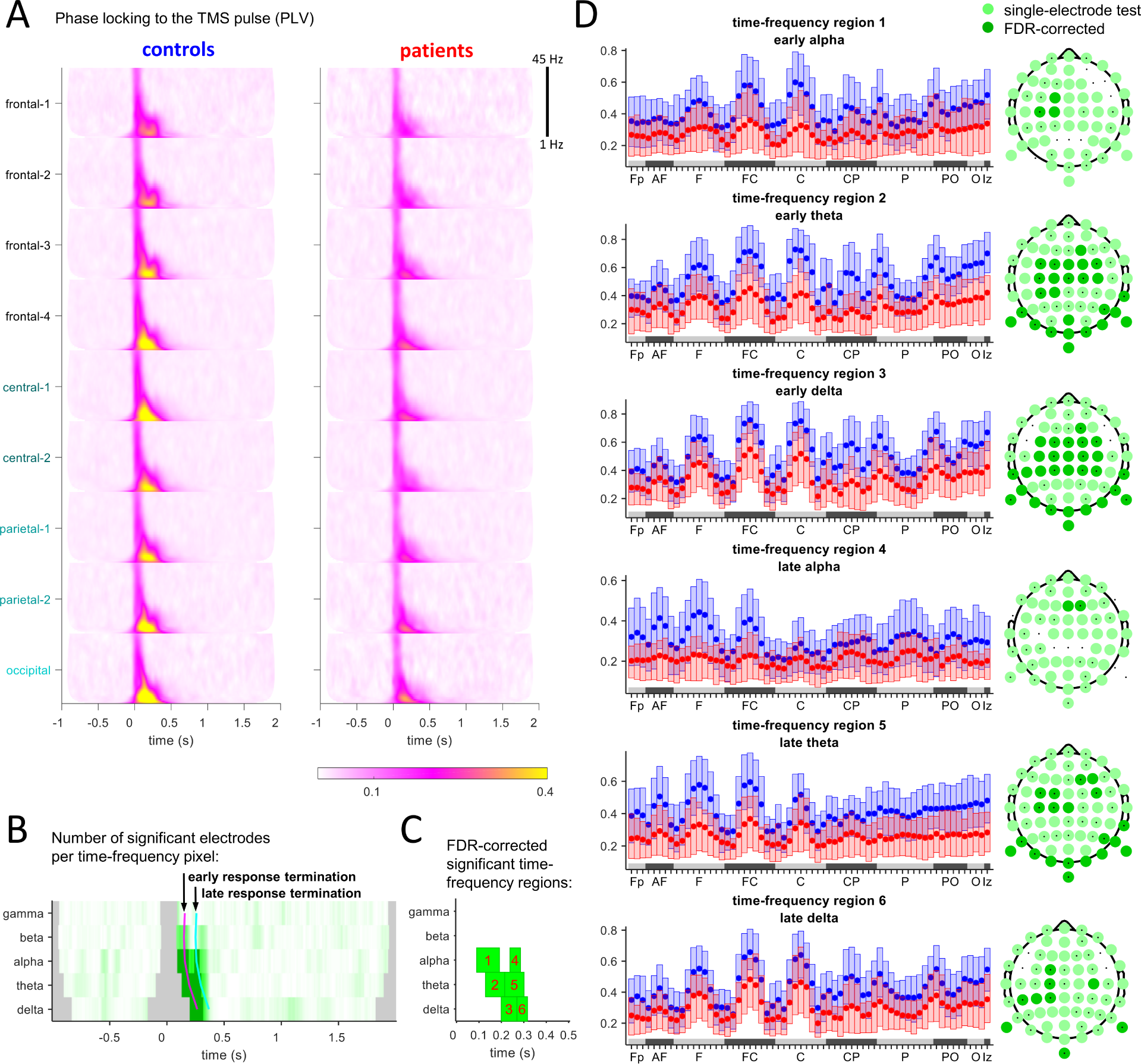
**A.** Phase locking value (PLV) for controls and for patients (displayed are averages of major electrode groups across trials). TMS was applied at time 0. **B.** For the PLV values at each time-frequency-electrode pixel, a statistical test was performed (p ≤ α=5 %). Shown is the number of electrodes that were significant at a time-frequency pixel. More intense green means larger number of significant electrodes. Dark green: Time-frequency pixels selected from those FDR-corrected time-frequency pixels for which there is at last one significant time-frequency-electrode pixel. Gray: excluded due to boundary and artifact interference. Early response: time span until 100 ms (magenta line), late: 100 ms to 200 ms (cyan line). **C.** Time-frequency regions obtained from grouping the dark green time-frequency pixels by frequency band and by response stage.

where the factor *GROUP* has the levels *Control* and *Patient*. The model has the fixed effects *γ*_00_, *γ*_10_ and the random effects *R*_*ij*_, *U*_0*i*_ with *R*_*ij*_ *∼ N* (0, *σ*), *U*_0*j*_ *∼ N* (0, *τ*_00_), where *N* (*a, b*) denotes a normal distribution with mean *a* and standard deviation *b*. The effects *R*_*ij*_ capture between-session (i.e. within-subject) variability. The effects *U*_0*j*_ encapsulate the between-subject variability and also incorporate the assumption of homogeneity of subject variance. Model fitting and significance testing was done with Matlab’s fitlme function. Notably, we repeated the (electrode-wise) statistical tests for Figs. 2A, 3B using the Wilcoxon rank sum test, which disregards the session structure. The results are very similar concerning the location of electrodes exhibiting a statistically significant difference between groups. The same testing procedure was also applied to the comparison of the alpha peak locations in Fig. 4B.

**Figure 4:**
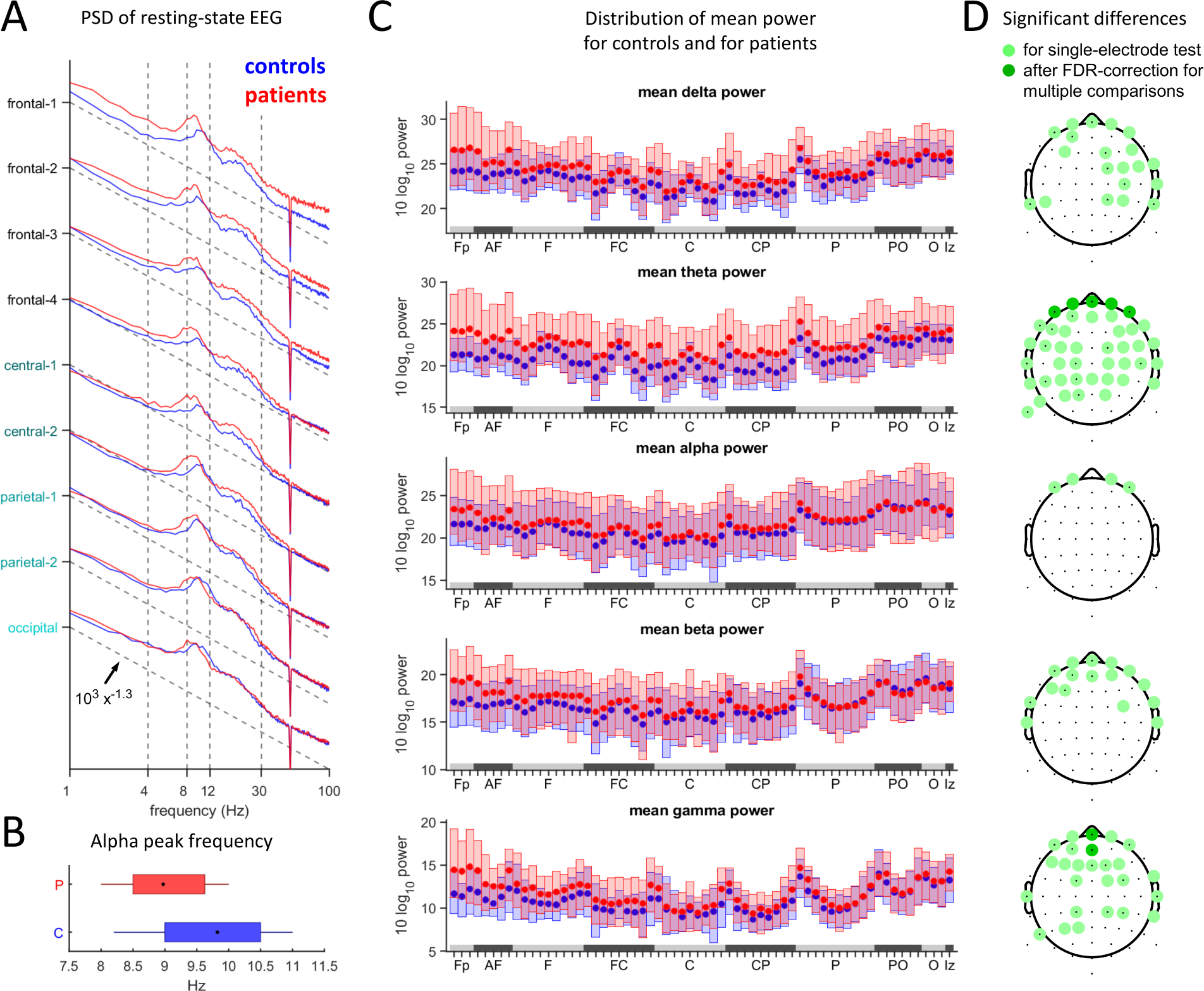
**A.** Log-log plots of resting-state EEG spectral power densities (PSDs) relative to a power law 10^3^x^−1.3^ (dotted line) for comparison. **B.** Alpha peak location for patients and controls. The black dot marks the mean, whiskers indicate the first and the third quartile. **C.** Boxplots of spectral power for the different frequency bands, for each electrode. **D.** Statistically significant group differences of spectral power in the different frequency bands. Tests were applied to each electrode separately (p ≤ α=5 %) and then corrected for multiple comparisons.

Following the separate testing, we corrected for multiple comparisons using the FDR procedure on the p-values of all time-frequency-electrode pixels together, yielding those time-frequency-electrode pixels that are significantly different.

Statistical tests for spectral power and quartile-based coefficients of variation computed from instantaneous amplitudes were done using the Wilcoxon test, because the amplitude distribution is strongly skewed and non-Gaussian. Consequently, we ignored the dependence of sessions of each subject and instead treated them as ‘independent subjects’.

### 2.7 Data visualization

Time-frequency pixels were further grouped into time-frequency regions for the purpose of visualization and interpretation of significant differences of the data (see Figs. 2A, B and Figs. 3B, C). To specify these groupings, we defined time windows of the TMS-evoked response by considering the time span from 25 ms to 100 ms as the early stage, the time span from 100 ms to 200 ms the late stage, and the time span from 200 ms onwards the very late stage. These time windows are depicted in Figs. 2A and 3B, C as magenta and cyan lines, respectively. The lines defining the time windows are curved because the Morlet wavelets are wider for lower than for higher frequencies. A time-frequency window was then defined as the intersection of a single standard frequency band and the time windows.

Time-frequency regions are then those areas within time-frequency windows for which there was at least one electrode whose time-frequency-electrode pixel was significant, as discovered by the mass univariate statistical analysis.

All boxplots were plotted such that the box edges correspond to the 20-th to 80-th percentile of the underlying data. The center dot indicates the mean.

For visualization purposes, major electrode groups were aggregated and averaged. This constituted the following electrode aggregates: frontal-1 (FP1, FPz, FP2), frontal-2 (AF7, AF3, AFz, AF4, AF8), frontal-3 (F7, F5, F3, F1, Fz, F2, F4, F6, F8), frontal-4 (FT7, FC5, FC3, FC1, FCz, FC2, FC4, FC6, FT8), central-1 (T7, C5, C3, C1, Cz, C2, C4, C6, T8), central-2 (TP7, CP5, CP3, CP1, CPz, CP2, CP4, CP6, TP8), parietal-1 (P9, P7, P5, P3, P1, Pz, P2, P4, P6, P8, P10), parietal-2 (PO7, PO3, POz, PO4, PO8), occipital (O1, Oz, O2, Iz).

## 3 Results

### 3.1 Power modulation due to the TMS pulse

Fig. 1 depicts the overall effect of the TMS pulse on the EEG amplitude. We computed time-frequency decompositions for each electrode and present the derived relative power in decibels, once as averages for the electrode aggregates (Fig. 1A, see Methods), and again as topographic plots of averages over consecutive time windows (Fig. 1B). This measures the power of the TMS-evoked response w.r.t. baseline power, and can hence be viewed as signal-to-noise ratio.

Three main qualitative effects are immediately apparent across the different time scales, ranging from tens of milliseconds to seconds and affecting several frequency bands. First, there is a reduction of the excitatory response in patients, which can be observed in two time-frequency regions. One is an immediate response to the TMS-pulse until around 300 ms in the delta, theta, and alpha bands (yellow and pinkish colors in Fig. 1B). The other appears in more central electrodes in the beta and gamma bands from around 500 ms to around 1 s (with white-pinkish in controls and white-bluish in patients, Fig. 1B).

The second effect is a facilitation of excitability, observable in two time-frequency regions. One is in the theta and alpha band, starting at around 300 ms and lasting at least until 1 s. The second occurs in the delta band after 700 ms until around 1.5 s. In controls, this effect appears as the bluish topographic plots in Fig. 1B, which are absent in patients.

The third effect we observe suggests a transition from a reduction in excitability in controls to facilitation of excitability in patients. This effect is visible in the time-frequency plots (mostly in the central-posterior-occipital electrodes), in the theta and alpha bands starting around 300 ms and lasting until around 1 s. It is also visible in the topographic plots that appear bluish (i.e. attenuated) in controls but have reddish (i.e. facilitated) additions for patients.

To quantify these three main effects, we performed an FDR statistical analysis of the time-frequency-electrode pixels (see Methods). This is presented in Fig. 2. Fig. 2A shows, at each time-frequency pixel, the number of electrodes that are significant within it. Fig. 2B shows the significant regions (see Methods) and these are visualized in more detail in Fig. 2C as boxplots and topographical plots. While the significant differences discovered in the statistical analysis result in smaller time-frequency regions than those that we observed qualitatively, the alterations of excitability are confirmed. Specifically, a reduction of excitability in the early response to TMS is apparent in time-frequency regions 1-3 at frontal, central, and posterior electrodes. An alteration of response excitability appears in time-frequency regions 4 and 5, where the mean power response electrodes turns from negative (power attenuation) in controls to positive (power facilitation) in patients. This effect is strongest at the posterior electrodes.

The differential alterations of excitability that we verified with TMS-EEG in our patient group thus include a reduced excitability in early responses and a facilitated excitability as well as a shift from attenuation to facilitation of excitability in later responses.

### 3.2 Phase locking to the TMS pulse

We used the phase-locking value (PLV) to assess phase angle similarity following the TMS-pulse at each electrode separately, across trials. The phases were derived from the time-frequency decompositions used for Fig. 1. Fig. 3A shows the PLVs for each frequency and for all electrode aggregates, indicating that PLVs are generally reduced in patients.

We performed a statistical analysis of the time-frequency-electrode pixels of PLVs (see Methods). The sum of all significant electrodes for each time-frequency pixel is shown in Fig. 3B. FDR-corrected time-frequency regions of significantly different time-frequency pixels are shown in Fig. 2C. Notably, these time-frequency regions are different from those regions that result from the power analysis of Fig. 2, and are significant in the early and late stages of the response. The PLVs at the delta, theta and alpha bands, mostly show significant differences at central and frontal electrodes, which are neighboring to the stimulation site, but also at posterior electrodes (Fig. 3D). We conclude that reduction of phase-locking to a stimulus can be verified with TMS-EEG in our patient group.

### 3.3 Spectral power in resting-state EEG

The resting-state EEG was also assessed for differences in power spectra and amplitude fluctuations. The power spectral densities (PSDs) averaged for all electrode aggregates are depicted in Fig. 4A, with several alterations noticeable. There is an increase of power in lower frequency bands in patients and generally an elevated power in more frontal electrodes. Furthermore, the location of the mean alpha peak is shifted towards lower frequencies (controls 9.8±0.9 Hz, patients 9.0±0.8 Hz). Both effects are statistically significant (Figs. 4B-D), with the session effects ignored in this case (see Methods).

We conclude that the resting-state EEG spectrum of the patient group exhibits alterations from control spectra, especially in the frontal electrodes. Decomposition into frequency bands suggests that in most electrodes these effects are due to the theta band, while in frontal locations they occur also in the delta, beta, and lower gamma bands. However, after correction for multiple comparisons, significant differences could only be verified in the theta band at prefrontal electrodes and in the gamma band in the central prefrontal-frontal electrodes.

### 3.4 Amplitude fluctuations in resting-state EEG

We next tested for alterations in amplitude fluctuations at each frequency. A simple possible marker is the amplitude variation. Because the amplitude mean, which relates to the spectral power, shows large differences between subjects, we normalize the variation by the mean. This quantity is called coefficient of variation (CV). Because the CV is sensitive to outliers, in our analysis we replaced it by the similar but more robust quartile-based coefficient of variation (qbCV). Fig. 5A shows the averaged amplitude medians and the corresponding qbCVs of the two subject groups averaged for different electrode aggregates. The qbCV of a pure white noise signal is denoted by *qbCV*_*Rayleigh*_ and is shown as a line. Deviations of the computed qbCVs from *qbCV*_*Rayleigh*_ indicate the presence of signals that are non-noise (see Methods). Clear differences in the qbCVs in all electrodes are evident between the groups. However, the set of electrodes and frequencies does not coincide with differences of the medians shown in Fig. 5B, demonstrating that the larger mean signal amplitude in patients is not generally associated with equally larger variation. This also indicates that the qbCVs and the medians probe two different aspects of the signal and of the noise. Boxplots for each electrode are shown in Fig. 5C, averaged over frequency bands. The qbCVs of controls are generally larger in the alpha and beta range, as is corroborated by statistical tests (Fig. 5D). Single-electrode statistical tests show differences in more than half of the electrodes in the alpha and beta bands. After controlling for multiple comparisons, 16 and 7 electrodes are different in the alpha and beta band, respectively, which are distributed in the frontal, central-parietal and posterior areas. The qbCVs of controls are not significantly different from those of patients in the gamma band, but are smaller in controls at a few frontal, parietal and posterior electrodes. In contrast, the delta band shows some minor differences, mainly at parietal and frontal-parietal electrodes, where the qbCVs are smaller in controls. The differences are significant in 3 electrodes.

**Figure 5:**
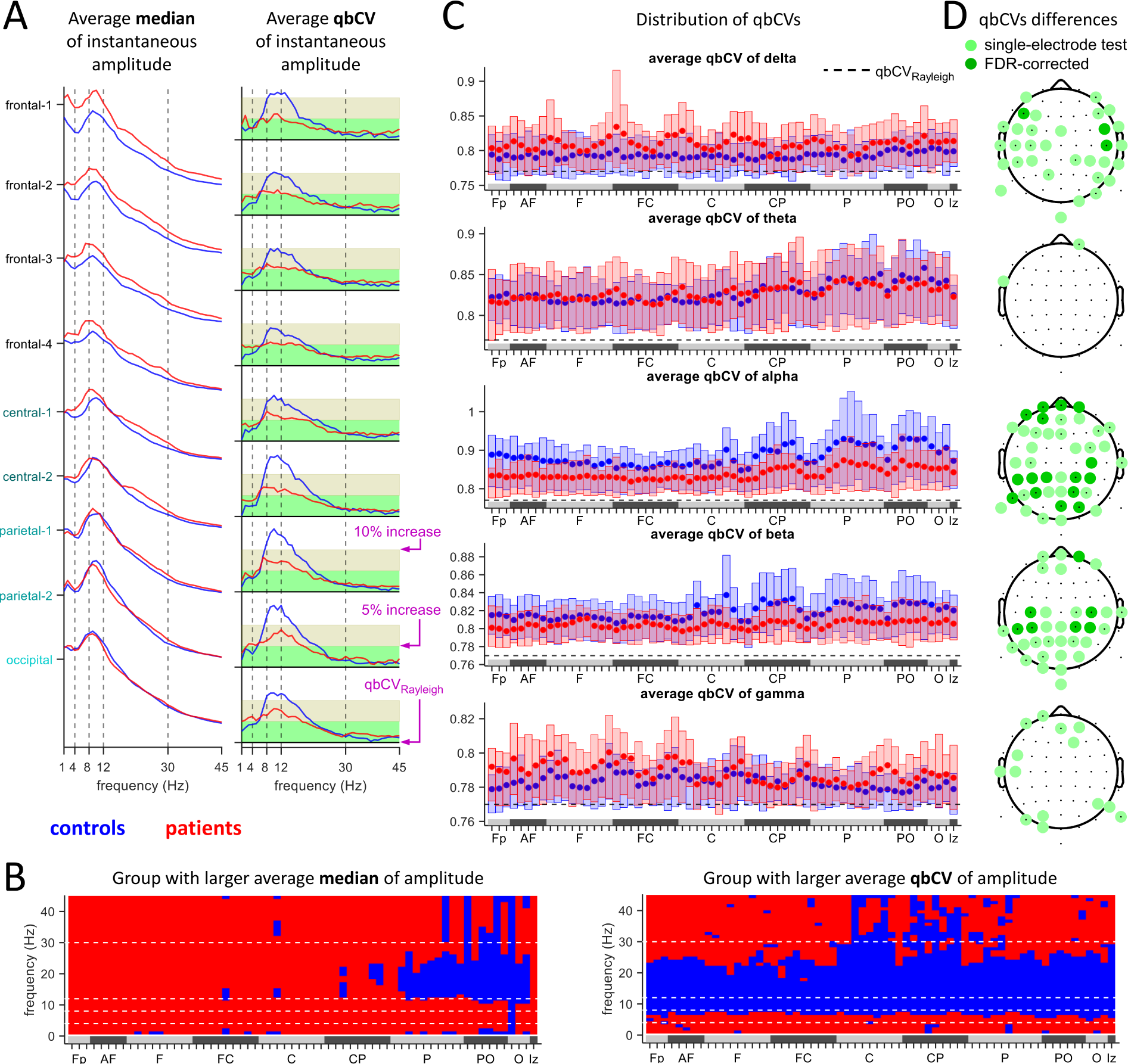
**A.** The averaged group median (left panel) and quartile-based coefficient of variation (qbCV, right panel) of the instantaneous amplitudes for different frequencies and electrode aggregates. The qbCV of amplitude fluctuations of a Gaussian noise signal qbCV_Rayleigh_ (and 5% and 10% increase of qbCV_Rayleigh_) is marked by horizontal lines. **B.** Color indication (patients: red, controls: blue), for each electrode and frequency, which group has the larger averaged group median (left) and qbCV (right). **C.** Boxplots of the qbCVs for the different frequency bands, for each electrode. The qbCV qbCV_Rayleigh_ of white-noise amplitude fluctuations is marked by the horizontal dashed line. **D.** Statistically significant group differences of qbCVs in the different frequency bands. Tests were applied to each electrode separately (p ≤ α=5 %) and then corrected for multiple comparisons.

We conclude that the resting-state EEG of the patient group exhibits a reduction of instantaneous amplitude fluctuations predominantly in the alpha and beta bands. In terms of the qbCV marker, the EEG in patients are closer to the value expected for random noise in the alpha and beta bands. Though a much smaller effect, the EEG of controls tends to be closer to the value of noise in the delta band.

## 4 Discussion

A hallmark of schizophrenia is the appearance of profound, global alterations of brain activity (Uhlhaas and Singer, 2010). The previously suggested noise hypothesis (Winterer et al., 2000; Rolls et al., 2008) on the nature of the phenomena underlying these alterations postulates the existence of inherent noise in brain activity in schizophrenia that degrades the brain’s ability to form stable spatial and temporal correlations between cells and cell populations. Stated in this form, the hypothesis can be evaluated for stimulus-evoked activity and for resting-state activity of the brain. Using TMS-EEG and resting-state EEG, we investigated various markers to evaluate the strength of coordinated population activity. Our analysis of the TMS-EEG confirms previously reported findings of evoked response properties based on power modulations and phase locking. In addition, we find clear differences of the resting-state EEG, which appear in the spectral power and the instantaneous amplitude. Using a fluctuation analysis of the instantaneous amplitude, we demonstrate that the schizophrenia resting-state data looks more similar to random noise than data from healthy controls. Overall, our results support the noise hypothesis in schizophrenia. In the context of the question to what extent intrinsic background activity in schizophrenia is more oscillatory than random, we provide support for the notion of more randomness in the EEG of schizophrenia patients. We point out that this randomness does not necessarily coincide with the inherent noise postulated by the noise hypothesis, however it may be indicative of its presence.

### 4.1 Alterations of evoked response processing

TMS application at the frontal-central site FC2 evoked strong responses lasting for more than 1 s. Our focus on spatial and temporal correlations and their relation to noise highlights that the power is modulated differently in the patient than in the control group, in different time windows and frequency bands (Fig. 1). While in controls there was increased power in various frequency bands in early time-frequency regions (Figs. 2A, B), in patients power was reduced. This contrasted later time-frequency regions, where patients exhibited increased power relative to the controls. This was clearly observable in the alpha and beta bands, but also qualitatively in the delta and theta bands (Figs. 1, 2A). However it was not significant in a more comprehensive statistical analysis.

The different time scales of the regions may be interpreted as two different response types to the TMS. The first type is a direct and local activation of the pulse, and the second subsumes all late-stage and long-range responses activated by the direct response (Ferrarelli et al., 2008; Frantseva et al., 2012; Radhu et al., 2015; Rogasch et al., 2014). In this picture, the changes of the signal-to-noise ratio in the different time-frequency regions (Fig. 2C) may have the following separate interpretations. At earlier response stages, there may be a reduction of excitability or an increase of inhibition (countering excitation), whereas at later stages there may be a reduction of inhibition (Frantseva et al., 2012) or facilitation of excitability (countering inhibition). An overall integration of these interpretations of these phenomena may be obtained in two different ways. One is that excitation and inhibition are both decreased, but the (global) excitation-inhibition balance is kept. The other is that the interplay between excitation and inhibition is generally different. The question which way is more likely may be considered based on the nature of the very late-stage response following the TMS-pulse (specifically, time-frequency regions 4 and 5 of Fig. 2C). If the relative power of the response in patients is viewed as oscillating around zero, it may be the result of reduced inhibition, favoring the first way. Conversely, if the relative power is viewed as positive, the inhibition in controls may have turned into excitation in patients, supporting the second way. Nonetheless, the late-stage response may only serve as a correlate, and an answer of the question requires more comprehensive evidence (Anticevic and Lisman, 2017; Selten et al., 2018).

These phenomena in the response to the TMS are consistent with the many known structural changes of brain tissue in schizophrenia. Indeed, alterations in connectivity have been reported (Krystal et al., 2017; Mikanmaa et al., 2019) during different developmental stages of the disease spectrum. Consistent with the excitability is a modification of local connectivity, such as a reduction of GABAergic cell density, myelination, connectivity, and alterations of gray matter density in progressed stages (Glahn et al., 2008). Consistent with alterations of signal propagation are abnormalities in long-range connectivity, such as changes of white-matter tracts (Tonnesen et al., 2018). In the context of TMS application, white matter integrity was demonstrated to directly influence how TMS-evoked activity spreads in the brain (Kearney-Ramos et al., 2018).

The power reduction of the TMS responses was accompanied by a reduction in locking of phase to the pulse, evaluated by the PLV. Significant group differences of PLV appeared in time-frequency regions at locations different from those for group differences of power. However, because both amplitude and PLV are related measures, this analysis is not independent of our findings regarding the changes in power. As expected, and reported previously for other paradigms (Winterer et al., 2000, 2004; Spencer et al., 2003), schizophrenia patients exhibit a reduction of phase locking. Indeed, we observe a significant reduction of PLV in the delta, theta, and alpha bands (Fig. 3). We note that the PLV can be artificially increased by volume conduction, which however would affect both groups to the same extent, and therefore these effects would work to reduce group differences rather than increase.

Our findings replicate previous reports for TMS-EEG in schizophrenia, such as Ferrarelli et al. (2008); Frantseva et al. (2012); Ferrarelli et al. (2012); Radhu et al. (2015); Ferrarelli et al. (2019); Rogasch et al. (2014); Shin et al. (2015) (see also Kaskie and Ferrarelli (2018)), and confirm previous observations in behavioral evoked-response paradigms (Winterer et al., 2000; Spencer et al., 2003). Alterations of longrange connections and inter-area synchronization in schizophrenia were reported in Hirvonen et al. (2017).

### 4.2 Alterations of resting-state power spectra

Various differences in the resting-state EEG spectra of schizophrenia patients were reported (Winterer et al., 2000; Uhlhaas and Singer, 2010; Goldstein et al., 2015), though inconsistencies in the literature have been pointed out (Hunt et al., 2017; Newson and Thiagarajan, 2019). In Ferrarelli et al. (2008), no significant differences in the spectra were found. In our dataset, we identified a significant shift of the alpha peak to lower frequencies (‘slowing’). This finding replicates Goldstein et al. (2015); Garakh et al. (2012); Yeum and Kang (2018); Murphy and Öngür (2019), and may be related to symptoms (Garakh et al., 2012) or duration of illness (Goldstein et al., 2015). When the significance of spectral power differences was evaluated electrode-wise, spectral power differences appeared only at frontal and parietal electrodes (Figs. 4A, D). The exception was in the theta band, which also showed differences in more central electrodes. After multiple-comparisons correction, all differences vanished except at two center-frontal electrodes in the gamma band and four prefrontal electrodes in the delta band. However, the significant difference in theta power may be an artifact of the alpha peak slowing together with our fixed definition of frequency bands.

### 4.3 Neuronal noise and alterations of resting-state amplitude fluctuations

Beyond the differences in the spectra, several other alterations of the resting-state EEG in schizophrenia have been reported. This includes analyses of microstates (Rieger et al., 2016; Michel and Koenig, 2018), long range temporal correlations (LRTC) (Nikulin et al., 2012; Sun et al., 2014), and amplitude excursions (Sun et al., 2014). While the origin of these differences is not known, they are consistent with the noise hypothesis. We quantified the variability of resting-state amplitude fluctuations at different frequency bands in a simple and straightforward way, using the qbCV for moving time windows of 40 s. Significant differences, corrected for multiple comparisons, appear in several frequency bands (Fig. 5), most notably in the alpha and beta bands. Both were also reported previously to exhibit significant reductions of LRTC in schizophrenia by Nikulin et al. (2012), whereas Moran et al. (2019) reports LRTC attenuations only for the beta band.

The qbCV is a measure similar to the coefficient of variation (CV), and can be viewed as evaluating the incidence and strength of random events, occurring when the instantaneous amplitude deviates from the amplitude average. These deviations are therefore indicative of a short-lived, oscillatory event. A larger qbCV can be considered as representing a richer repertoire of events and hence of cortical activity, although it formally equates to a higher noise-to-signal ratio. Support for this view comes from experimental measurements of the spiking activity in rodents (de Vasconcelos et al., 2017; Fontenele et al., 2019), in which the CV, computed on the timescale of tens of seconds, was used to characterize different cortical states.

A second interpretation of the qbCV that goes beyond the mere detection of group differences is that a signal with a qbCV that is closer to *qbCV*_*Rayleigh*_ (see Methods) is indicative of increased randomness. However, this interpretation is not as straightforward and relies on the following additional assumptions.

The first assumption regards our model of the narrow-band filtered EEG signals, which we assume to consist of two parts. One part comprises the modulated oscillatory events and consists of oscillations at frequencies around the center frequency of the band. The amplitude envelope of this part can vary slowly compared to the center frequency. The other part comprises of white noise with a constant strength. Hence, as the amplitudes of the event occurrences tend to zero, the qbCV of the instantaneous amplitude approaches *qbCV*_*Rayleigh*_.

The second assumption regards the relationship between amplitude and duration of the oscillatory event occurrences. Generally, the shorter the coverage by oscillatory events and the larger the event amplitudes, the larger is the qbCV. Furthermore, longer coverage by oscillatory events can make the qbCV very small, and even smaller than *qbCV*_*Rayleigh*_ if the event amplitudes are large enough (see the examples discussed in Methods). So the interpretation that resting-state EEG activity is closer to noise is only true if we exclude that the majority of patients had rather long and rather strong occurrences of oscillatory events that are fine-tuned such that their qbCVs are close to *qbCV*_*Rayleigh*_. In other words, this interpretation relies on the reasonable assumption that such very specific fine-tuning is unlikely.

Despite the generally increased mean signal amplitude in patients, the signal variation is not generally increased to the same extent. This manifests itself as a striking reduction of qbCVs in the alpha and beta bands compared to controls. Given our assumptions, we interpret these smaller qbCVs indicate a lower signal content in the alpha and beta band. Furthermore, differences in the qbCVs in the delta band are indicative of different activity patterns in this band.

### 4.4 Specificity of the investigated features for schizophrenia

Classification studies of intrinsic activity and evoked potentials in schizophrenia and psychosis indicate that achieving a comprehensive explanation that links alterations of intrinsic activity and evoked potentials would be difficult (Hudgens-Haney et al., 2017; Rolls et al., 2017). In fact, Clementz et al. (2017) were constituted three different ‘biotypes’ from a battery of properties, which do not coincide with clinical classification schemes. Hudgens-Haney et al. (2017) found alterations, corresponding to the biotypes, in EEG background activity and in certain evoked potentials. Although the origin of these phenomena is not clear, it is consistent with the large overlaps between the healthy range and the patient range in our measurements (see e.g. the boxplots in Figs. 2-5).

To conclude, our results corroborate and expand the previously reported evidence for the noise hypothesis in schizophrenia, obtained by investigating evoked potentials and resting state data. In TMS-EEG, we describe and confirm various alterations of evoked potentials. In resting-state EEG, we confirm alterations of amplitude fluctuations schizophrenia, using a new analysis based on a simple ratio measure of dispersion and mean. We suggest to distinguish between the extent of abnormal randomness and of abnormal oscillations in the brain activity, and note that this generalizes to both evoked potentials and resting-state EEG. A reasonable interpretation of our findings supports the possibility that the schizophrenia brain is characterized more by abnormal randomness.

### 4.5 Limitations

Our study has several limitations. One results from the placement of the TMS coil. The coil position was determined using manual measurements and a customized stand, rather than with neuro-navigation methods. This has likely resulted in small deviations of coil placement between sessions. A second limitation results from the fixed stimulation strength at 80 % maximal stimulation strength rather than an adaption to motor threshold. Both of these flaws may have increased inter- and intra-subject variability of the EEG measurements. However, we expect these variations to be distributed similarly in the two groups and not to be biased towards one group. In this case, the increased variability works against significance of the measured effects, and hence may increase the robustness of our findings. Moreover, despite these expected deviations, our results are robust across sessions, thus reducing the possibility that our findings are merely the result of a bias in coil placement or stimulation strength.

A third limitation of our study concerns the fact that our patients were medicated. Antipsychotics are known to generate EEG abnormalities (Centorrino et al., 2002).

A fourth limitation concerns the caffeine and nicotine intake of the subjects. We asked subjects to keep the intake (as well as sleep) consistent on experiment day, but since nicotine and caffeine consumption is typically increased in schizophrenia patients, it cannot be excluded that our EEG measurements may be altered (Bchir et al., 2006) Lindgren et al. (1999). For example, such intake can increase the alpha peak location.

